# Emergent properties of coupled bistable switches

**DOI:** 10.1101/2021.06.15.448553

**Authors:** Kishore Hari, Pradyumna Harlapur, Aditi Gopalan, Varun Ullanat, Atchuta Srinivas Duddu, Mohit Kumar Jolly

## Abstract

Understanding the dynamical hallmarks of network motifs is one of the fundamental aspects of systems biology. Positive feedback loops constituting one or two nodes – self-activation, toggle switch, and double activation loops – are commonly observed motifs in regulatory networks underlying cell-fate decision systems. Their individual dynamics are well-studied; they are capable of exhibiting bistability. However, studies across various biological systems suggest that such positive feedback loops are interconnected with one another, and design principles of coupled bistable motifs remain unclear. We wanted to ask what happens to bistability or multistability traits and the phenotypic space (collection of phenotypes exhibited by a system) due to the couplings. In this study, we explore a set of such interactions using discrete and continuous simulation methods. Our results suggest that couplings that do not connect the bistable switches in a way that contradicts the connections within individual bistable switches lead to a steady state space that is strictly a subset of the set of possible combinations of steady states of bistable switches. Furthermore, adding direct and indirect self-activations to these coupled networks can increase the frequency of multistability. Thus, our observations reveal specific dynamical traits exhibited by various coupled bistable motifs.

## Introduction

Multistability in biological networks is a crucial phenomenon responsible for various functions such as reversible switching between cell states and consequent phenotypic heterogeneity [1–5]. For non-linear systems such as biological networks, a necessary condition for multistability is the existence of positive feedback loops [6,7]. A feedback loop is defined as a series of connected edges in a directed regulatory network that starts and ends at the same node. Positive feedback loops are defined as feedback loops with an even number of inhibitory links [8]. Thus, in simple one-node and two-node networks, self-activation, a mutual activation between two nodes, and a mutual inhibition between the two nodes are examples of positive feedback loops [9,10]. Understanding the dynamics of such feedback loops and their interconnections in a large regulatory network can help decode cellular decision-making.

The simplest of them, a self-activation, involves a single node enhancing its own production/ accumulation directly or indirectly and can give rise to two states, one with low expression of the node (“OFF” or 0) and the other with high expression (“ON” or 1) [11]. Mutual inhibition between two nodes, also known as a toggle switch (TS), is among the most commonly observed motifs in systems involving cellular differentiation in development and disease [12– 14]. A TS involves two biomolecules inhibiting the production/accumulation of each other, leading to two states (or phenotypes) in the system, wherein each phenotype, levels of one of the nodes is high (“ON”) while the other is comparatively low (“OFF”) – (1, 0) and (0, 1) [15]. The addition of one or more self-activation to a TS has been shown to give rise to hybrid states where the levels of both nodes involved in a TS are at comparable levels, often intermediate to those seen in the two canonical states seen in a TS, i.e. (1, 1) or (½, ½) state [12,16]. For instance, in epithelial-mesenchymal transition (EMT), where multiple TS have been reported between transcription factors and microRNAs, such hybrid E/M states are witnessed extensively [17–19]. Further, toggle switches with self-activation (TSSA) are seen at cell decision-making branch-points in embryonic development, too [12]. Similar to a TS, a double activation (DA) network can also bring about bistability, but the two stable states emerging from a DA are non-overlapping with those seen from a TS – (0, 0) and (1, 1) [20].

Biological networks often contain coupled DA and TS, which have been investigated on a case-to-case basis [20–26]. A bidirectional coupling between two toggle switches – for instance, one regulating EMT and another affecting stemness – can help coordinate the behavior enabled by individual bistable motif [27]. In a more generic scenario of such cases, a question often arises whether all possible combinations of steady states allowed by both individual TS or DA motifs can indeed be observed and whether any new states can emerge depending on the extent of strength and signs of different coupling links. Here, we address this question in the context of various instances of couplings between the simplest positive feedback loops discussed above: toggle switch (TS), double activation (DA), and self-activation.

## Materials and Methods

### Network simulation: Random Circuit Perturbation (RACIPE)

RACIPE [28] is a computational method that takes in network topology as the input and generates a set of corresponding Ordinary Differential Equations (ODEs) with randomized kinetic parameters to represent the network dynamics. A node T in the network having P_i_ and N_j_ activating and inhibiting nodes, respectively, acting on T, the ODE representing the node will be given by:

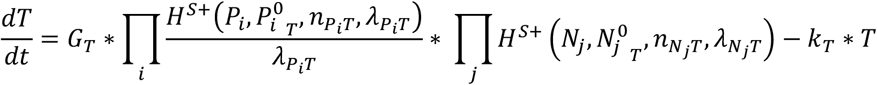

Where T, P_i_, and N_j_ represent the concentrations of the species, G_T_ the production rate, k_T_ the degradation rate of the node T. P_i_ ^0^_T_ and N_j_ ^0^_T_ are the threshold concentrations of T required for the activation of the respective nodes. *λ* represents the fold change in T’s expression upon being activated or inhibited by the individual input nodes. The Shifted-Hill Equation, H^S^ represents the regulation of T by the input nodes.

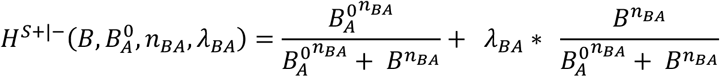

RACIPE generates multiple parameter sets by randomly sampling from predefined ranges for specific parameters as listed in the table below:

| Parameters | Minimum Value | Maximum Value |
| --- | --- | --- |
| Production Rate (G) | 1 | 100 |
| Degradation Rate (k) | 0.1 | 1 |
| Fold Change (Inhibition $\lambda^-$) | 0.01 | 1 |
| Fold Change (Activation $\lambda^+$) | 1 | 100 |
| Hill Coefficient (n) | 1 | 6 |
| Threshold | The range depends on inward regulations—half functional rule. |  |

After the parameter sets are generated, RACIPE solves the ODEs with multiple initial conditions, using Euler’s method, to obtain the steady states for a corresponding parameter set, which is then written in the output files. The RACIPE code used for simulating the networks can be found at https://github.com/simonhb1990/RACIPE-1.0. Simulation of all the networks was done in triplicates, with each replicate having 10000 parameter sets (sampled as a uniform distribution from their respective ranges) and 100 initial conditions, the results of which were compiled and analyzed using custom Python3 scripts.

### Calculation of Steady state Frequencies from RACIPE data

The steady state expression levels for all the nodes of the network were normalized using the following equations:

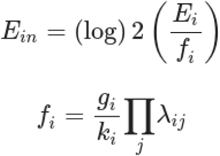

Where, Ein is the normalized steady state expression, E_i_ the steady state expression level, f_i_ the normalizing factor, g_i_ the production rate and k_i_ the degradation rate of the ith node corresponding to the current steady state. *λ*_ij_ are fold change values on ith node due to the input nodes j.

The normalized expression levels were then converted to z-scores according to the below equation:

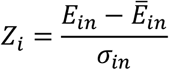

Where Z_i_ is the z-score of the expression level of the ith node, E_in_ the normalized expression level of the ith node, 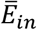 the combined mean and *σ*_*in*_ the combined variance.

The z-scores are then “booleanised” into 1’s and 0’s representing high or low expression levels which are concatenated to represent the steady state for a particular parameter set. The frequency of the occurrence of a particular steady state is calculated by counting its occurrence in all the parameter sets.

### Link strength analysis of RACIPE parameter sets and probability of occurrence of steady states

The probability of obtaining each state in a given network is computed by considering link strength data for each parameter set. For a given parameter set, the strength of a link between node i and node j is calculated as follows:

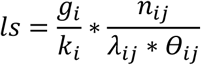

Where *g*_*i*_ is the production rate of node i, *k*_*i*_ is the degradation rate, *n*_*ij*_ is Hill coefficient, *λ*_*ij*_ is the fold change in the production rate of j due to i and *θ*_*ij*_ is the threshold parameter for the link. A threshold is defined for each link, which is the mean link strength of that link across all parameter sets. For each parameter set, the number of links that are active is calculated as the links that have their strength higher than the corresponding threshold. Then, a probability score is assigned for each parameter set, equal to 0.5^n, where n is the number of links that are active for that parameter set. The probability of a steady state occurring is then calculated, overall parameter sets that give rise to that particular steady state, is given by :

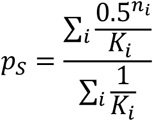

Where K_i_ is the number of steady states the i^th^ parameter has.

### Network simulation: Boolean formalism

We obtained steady state distributions for our network using Boolean modeling with Ising Model formalism and synchronous or asynchronous update algorithms. Each state is represented with the Boolean function of N elements, where N is the number of nodes in the network. The state of the system varies over time through discrete transitions, eventually stabilizing in an attractor, starting from a randomly selected set of initial conditions. The nodes are updated according to the following equation:

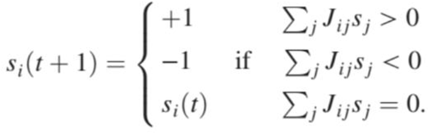

where s_i_ (t) denotes the expression levels of node *i* at time *t*. s_i_ = +1 means that the node is highly expressed (i.e., ‘ON’ state), otherwise s_i_ = -1. J depicts the interaction matrix of the network. J_ij_ = 1 indicates that node *i* promotes the expression of node *j*, J_ij_ = --1 implies that node *i* inhibits the expression of node *j*, J_ij_ = 0 implies no regulatory interaction from node *i* to node *j*.

The packages used to simulate the networks using ising formalism are: https://github.com/ComplexityBiosystems/bmodel and https://github.com/csbBSSE/bmodel_julia.

### Boolean state-transition graph

State transition graphs [29] are generated by tracking all possible steady states that each initial condition can converge to. These initial-final state pairs are then used to construct the edges in the transition graph. The representation of these graphs is obtained using the package *igraph* in R 4.0.

### Jensen-Shannon Divergence (JSD)

Jensen-Shannon Divergence [30] was used to quantify and compare the differences in steady state distributions of RACIPE and Boolean simulations. JSD for any two discrete frequency distributions P(x) and Q(x) is calculated by:

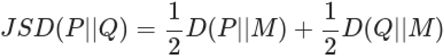

Where M = ½(P+Q) and D is the Kullback-Leibler Divergence given by:

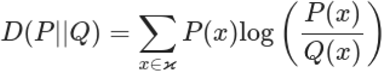

When base 2 logarithm is used for JSD calculations, the score varies from 0 to 1, with 0 indicating identical distributions and 1 indicating the two distributions have no overlap. The function *jsd* from the package *philentropy* in R 4.0 was used to calculate JSD.

### Statistical analysis

All error bars given in figures represent mean ± sd where sd is the standard deviation.

### Programming languages and codes

The analysis was done using custom codes written in *Python 3*.*6* and *R 4*.*0*. All plots were generated using *ggplot2* package of *R 4*.*0* except for state transition graphs, for which the *igraph* package was used. The codes required to reproduce the analysis are given in https://github.com/csbBSSE/CoupledBistableSwitches.

## Results

### Phenotypic distribution and multistability seen in bistable switches

To analyze the effects of coupling of TS and DA, we chose to simulate the networks using discrete (Boolean-Ising) and continuous (RACIPE) approaches. The former, as the name suggests, considers binary levels of expression levels (−1 = OFF, 1= ON) for each node and assigns equal weights to regulatory edges incumbent upon a given node (−1 for inhibition and 1 for activation) [31]. This approach is independent of any kinetic parameters and offers a good qualitative insight into the steady-state behavior [32]. On the other hand, the later formalism, RACIPE, considers a set of coupled ODEs representing the network dynamics and samples the kinetic parameters for the set of ODEs from a predefined n-dimensional parameter space (n = N*2 + E*3, where N is the number of nodes and E is the number of edges in the network) and simulates network dynamics for sampled parameter sets across diverse initial conditions [28]. With sufficient sampling for parameter sets and initial conditions, we can obtain an ensemble of network behavior that is representative of the network behavior in a wider parameter regime. For ease of comparison with Boolean, we discretized the steady state levels obtained from RACIPE (see **Methods** section for more details on these formalisms).

As a first step, using both these formalisms, we analyzed the TS and DA motifs to understand their steady-state and multistability patterns. The TS motif displays two major steady states in which either of the genes is expressed in higher concentrations while the other is repressed (i.e., 01 or 10) (**Fig 1A**). This trend was seen in both RACIPE and Boolean results, with Boolean showing a slightly higher frequency of these states (∼50% each in Boolean vs. 41% and 44% in RACIPE). RACIPE also enabled states in which both the genes were expressed in high concentrations (i.e., 11), albeit at a low frequency (∼15%). This behavior is expected because RACIPE can sample parameter combinations that might render the inhibitory links weak, thereby allowing steady states with both genes having high expression levels. The differences between RACIPE and Boolean, however, are relatively small, as quantified by Jensen-Shannon Divergence (JSD), a metric that measures the difference between two given frequency distributions [30]. The value of this metric varies between 0 and 1, with 0 indicating identical distributions and 1 indicating non-overlapping distributions. JSD metric, for this case, is 0.08 (**Fig 1A**).

**Fig 1.**
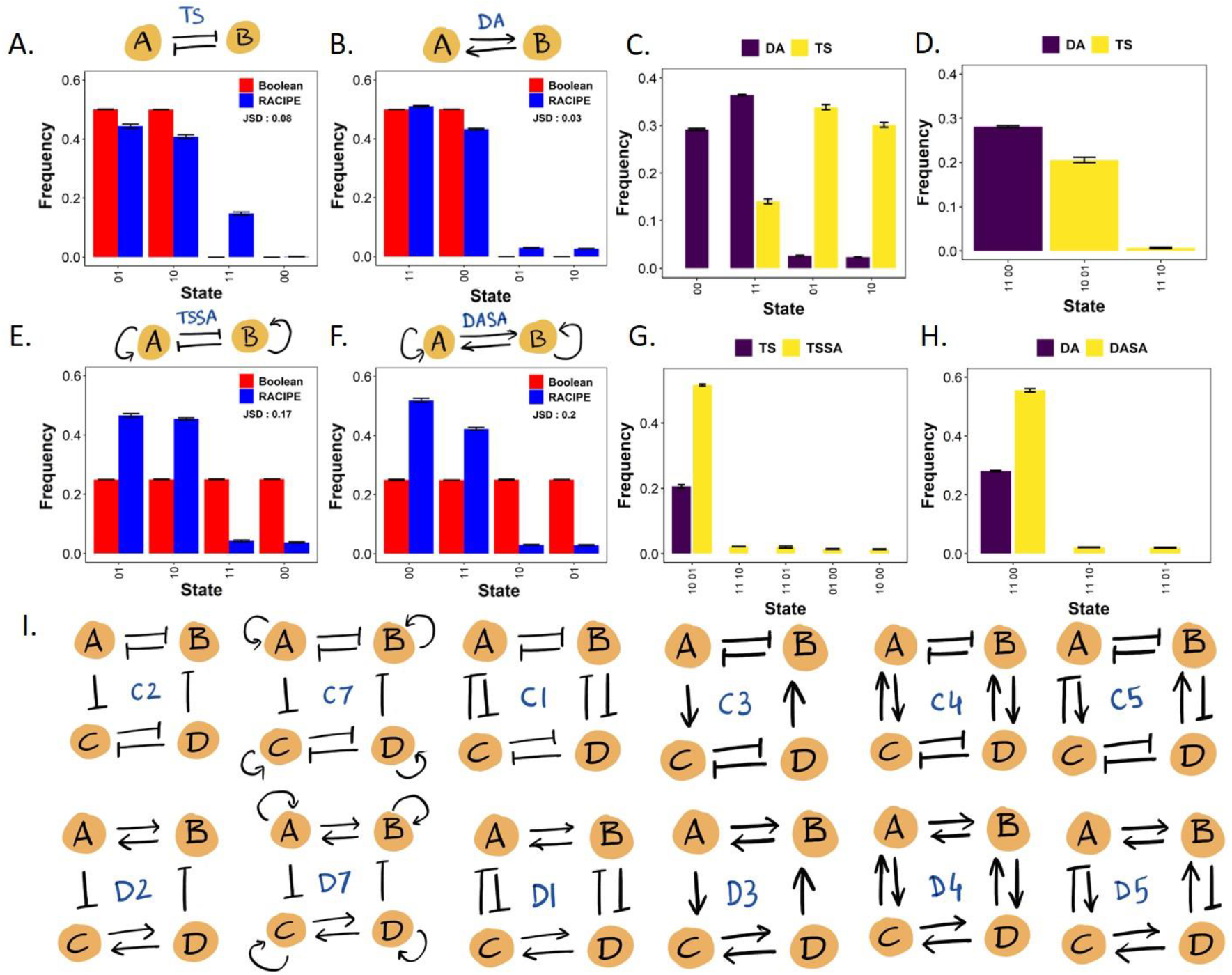
Phenotypic composition and multistability of bistable switches. A) Phenotypic distribution of TS motif. B) Phenotypic distribution of DA motif. C) Comparison of monostable steady states of TS and DA motifs. D) Distribution of multistable steady states of TA and DA. E) Phenotypic distribution of TS motif with self-activation at each node (TSSA). F) Phenotypic distribution of DA motif with self-activation at each node (DASA). G) Comparison of multistable steady states of TS and TSSA. H) Comparison of multistable steady states of DA and DASA. I) List of coupled networks analyzed.

The DA loop predominantly converged to two steady states (∼50% each in Boolean vs. 42% and 50% in RACIPE) in which both the nodes showed the same expression levels – 0 or 1; i.e., the states 11 or 00 (**Fig 1B**). Similar to the trends seen for TS, RACIPE and Boolean yielded consistent results (JSD = 0.03) with the two additional states (01, 10) identified by RACIPE having a low frequency (∼4% each). Thus, the behavior of TS and DA loops is largely non-overlapping: merely a 4-5% frequency of the most dominant TS states (10, 01) is seen in DA and *vice-versa* (frequency of 00, 11 states in a TS) as per RACIPE simulations.

We then analyzed the “phases” of the motifs given by RACIPE. A phase is defined as a combination of one or more co-existing steady states for a given parameter set. A parameter set for which the system can converge to only one steady state, irrespective of the initial conditions, is said to belong to the corresponding monostable phase. Similarly, a parameter set for which convergence to more than one steady state is observed when starting from an ensemble of initial conditions is said to be multistable. We calculated the frequency of different monostable parameter sets/phases and observed that their frequency distribution largely resembles (**Fig 1C**) the frequency of all states seen earlier (**Fig 1A-B**). Furthermore, the multistable phases of both TS and DA were composed of the combination of their most frequent monostable steady states – {10, 01} for TS and {00, 11} for DA (**Fig 1D**).

A variation of TS and DA motifs that is commonly observed in biological networks is the presence of self-activation on one or both of the nodes [16]. Thus, we investigated the dynamics of TS and DA with both the nodes directly self-activating themselves – TSSA (toggle switch with self-activation), DASA (double activation with self-activation). While both TS and DA display characteristic states in Boolean simulations, the addition of self-activation neutralizes this trend by enabling equal frequency for all four possible states – 00, 01, 10, 11 – for both the circuits (**Fig 1E, F**). RACIPE simulations, however, still result in the dominance of the characteristic states – 10 and 01 for TS, 00 and 11 for RACIPE. In addition, RACIPE simulations indicate that the DA and TS motifs with self-activation have a higher frequency of multistable phases than the ones without it (**Fig 1G, H**). TSSA and DASA also showed tristable phases (**Fig S1 B, D**), endorsing previous observations that multistability is enhanced in networks with more positive feedback loops (TSSA or DASA has two more such loops than TS or DA correspondingly) [7]. Similar to TS and DA, bistable phases are composed of the most frequent states observed in phenotypic distributions and those in the monostable phases (**Fig S1 A, C**).

Having analyzed the dynamics of individual TS and DA motifs, we moved on to investigating the dynamics of various coupled TS and DA motifs. We simulated each of the coupled networks (**Fig 1I, S1E**) using RACIPE and Boolean formalisms and analyzed their phenotypic distributions, multistability, and the effect of adding self-activation, similar to our examination of DS and TA. We expect that the outcome of coupling will be a combination of the nature of the individual constituents (TS or DA) and the nature (direction and sign) of the coupling link(s).

### Coupling bistable motifs through activation and inhibition links

We connected the bistable motifs using different inhibition/activation links and feedback loops. The first among those couplings are bistable switches connected via two inhibition links, one going from the TS between A and B to that between C and D and one in the reverse direction. The phenotypic distribution for this circuit (C2) where A inhibits C and D inhibits B is shown in **Fig 2A**. For this circuit, the most dominant states according to both Boolean and RACIPE are (A, B, C, D) = (1, 0, 0, 1) and (0, 1, 1, 0) (Boolean: ∼33% each, RACIPE: ∼38% each). In these states, the pairs of nodes involved in individual TS – (A, B) and (C, D) – have opposite levels of activity. Similarly, the two nodes across the two TSs connected via inhibitory links – (A, C) and (B, D) – show opposite levels of activity. Because A inhibits C, which inhibits D, and D inhibits B, which inhibits A, A and D effectively activate each other in this network; thus, A and D show the same levels of activity in both the dominant states – similar to what would be expected between a DA formed by A and D.

**Fig 2.**
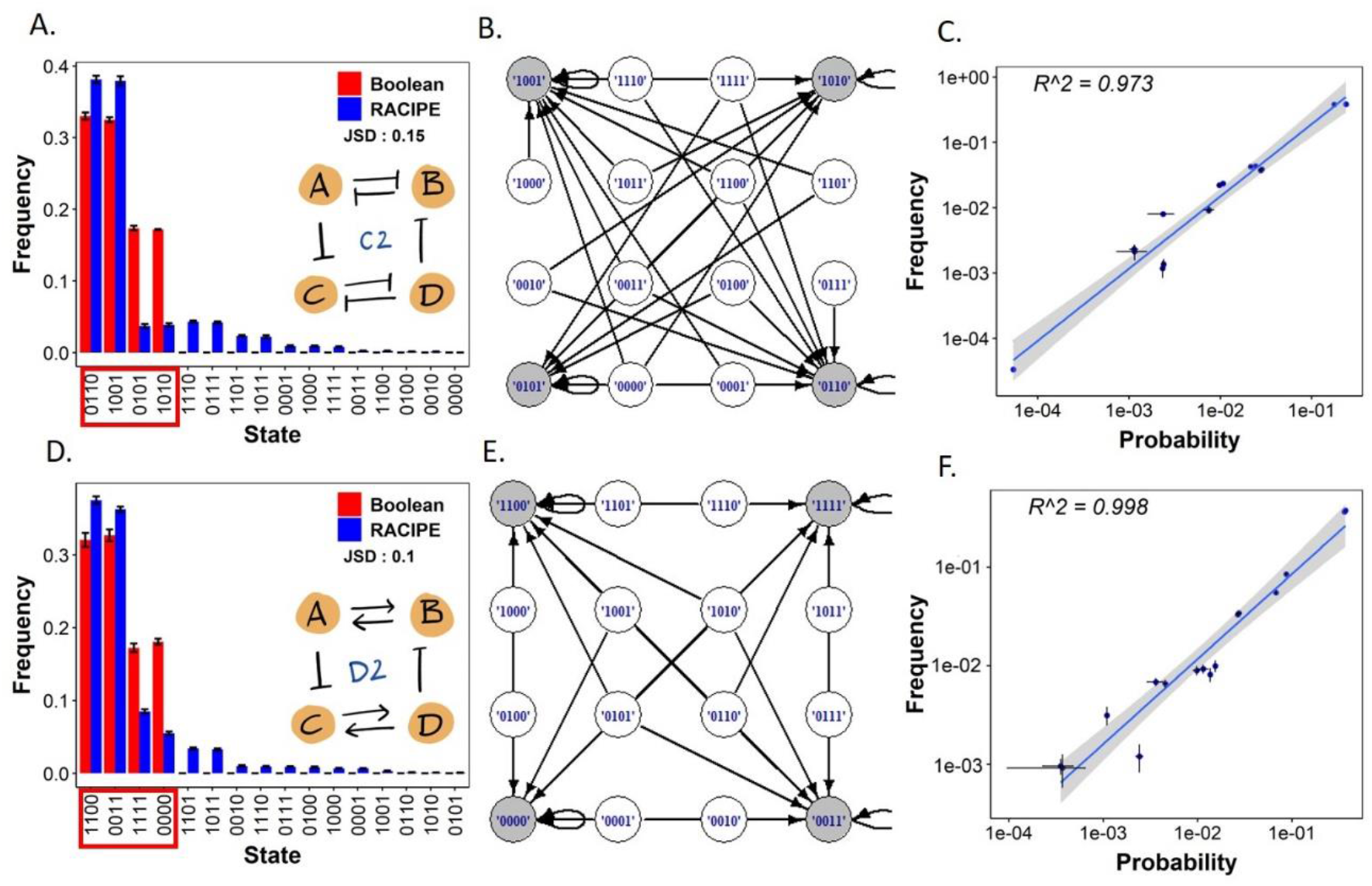
Coupling bistable switches with two inhibitory links. A) Steady state distribution of C2; states reported by Boolean simulations are highlighted in the red box. B) State transition graph of C2. C) Link-strength analysis of parameter sets of C2. X-axis represents the probability of a state occurring calculated from the link strengths, and the y-axis is the frequency of the steady states obtained from RACIPE. The R-squared value of the linear fit is indicated in the plots. D-F) Same as A-C but for D2.

In C2, two other states have considerable frequency (∼17% each) in Boolean results: 1010 and 0101. These two states still maintain the nature of individual toggle switches but fail to satisfy the bidirectional couplings, leading to a reduced frequency. The state transition graph (STG) (**Fig 2B**) depicts the basins of attraction corresponding to the four stable states. The graph was constructed by identifying all possible, stable steady states to which each of the 16 possible initial conditions can converge to via an asynchronous Boolean update. Due to asynchronous Boolean update, each state can potentially converge to multiple stable states [33]. For instance, the state 1100 can converge to all four stable states of the network, as seen in the STG. Empirically speaking, the states closer to 1001 and 0110 (different from the stable states by a single node, 0111, 1000 for example) always converge onto these states, while those closer to 1010 and 0101 can converge to either of the four states, thus indicating a wider basin of attraction (number of states converging into them) for 1001 and 0110 states.

In contrast to Boolean results, RACIPE shows a stronger dominance of 1001 and 0110 states for C2. While 1001 and 0110 only have a two-fold higher frequency than 1010 and 0101 in Boolean, the corresponding frequency of these states in RACIPE is >9 fold higher than that of any other state (∼38% vs. ∼4%). This trend was unexpected because RACIPE simulates a sample of parameters from a large parameter space and thus is likely to allow for more diversity in terms of phenotypes seen. While we see more phenotypes in RACIPE, the frequencies of these phenotypes follow a different distribution than that seen in Boolean. We hypothesize that the reason for this different behavior could arise from RACIPE sampling parameters in such a way that each link has only a 50% chance of being active (“half-functional rule”) [28]. Hence, the minimum number of links that need to be functionally active (“ON”) for achieving a state could dictate these frequency discrepancies. To test this hypothesis, we calculated the number of edges that have to be “on” for an ensemble of parameter sets to converge to a particular state. This calculation used the link strength metric we defined previously [34], and we found that the (expected) probability of a steady state being achieved as calculated has a strong linear relationship with the corresponding steady state frequency as obtained from RACIPE simulations (**Fig 2C**, R^2^ = 0.973 for a linear fit).

Next, we coupled toggle switches with activation links instead of inhibition links (C3, i.e., A activates C and D activates B). The results obtained showed adherence to similar rules as those seen in C2 (**Fig S2A-C**). The top two states – 0101 and 1010 (∼33% frequency each in Boolean and ∼38% frequency each in RACIPE; same as C2) – follow the individual toggle switch restrictions (i.e., A and B have opposite levels, and so do C and D) and the coupling configurations (A and C are both consistent with one another, and so are B and D). While Boolean simulations show two more states (1100 and 0011) with a frequency of ∼17% each, all of the states seen in RACIPE except 0101 and 1010 have frequencies less than 5%. The corresponding STGs show that the neighboring states of 1010 and 0101 converge to 1010 and 0101, but those of 1001 and 0110 do not converge to 1001 or 0110, thus potentially explaining their lower frequency. Similar to the earlier case, RACIPE results can be explained by the half-functional rule of RACIPE **(Fig S2C)**.

After analyzing the coupling of two TS, we characterized the state-space of coupling between two DA motifs by single inhibition in either direction, i.e., A inhibits C and D inhibits B (D2). The top states seen in Boolean simulations were 0011 and 1100 (∼32% frequency each), where only one of the two DA switches is “active” (**Fig 2D**). This observation can be explained by considering the DA motif as an effective one-node because both the nodes in a DA have the same state – 0 or 1 – thus, 00 and 11 are the most dominant states. Therefore, the inhibitions connecting the two DA loops can be construed as a toggle-switch-like behavior. However, the two states seen (∼18% frequency each) in Boolean simulations have both DAs acting together (1111 and 0000), which are not compliant as per a toggle switch-like configuration. Nevertheless, these states do not contradict the simplification of DA acting as a single unit. However, comparing this behavior with that of TSSA suggests that the D2 network can be simplified as a TSSA instead of a TS. One limitation of this simplification that comes in Boolean results is that while TSSA allows equal (∼25%) frequency of all four possible states – 0011, 1100, 0000, and 1111 (**Fig 1E**), the frequency of these states obtained by simulating D2 are not equal, with 0011 and 1100 being almost twice as more frequent than 0000 and 1111 (∼33% vs. ∼17%; **Fig 2D**).

Similar to C2 and C3, the dominant steady states of D2 also have their neighbors exclusively in their basin of attraction, while the less frequent ones share their neighbor states with the top two states. We also calculated the probabilities for the steady states using link strength analysis, similar to **Fig 2C** (**Fig 2F**). The state probabilities have a strong linear relationship with the RACIPE frequency values. Interestingly for D2, probabilities, and RACIPE frequency values are almost equal to each other (slope of linear fit ∼=1, intercept ∼=0). Similar results have been observed when the DAs were connected by activation links instead of inhibition ones, which can be explained by replacing the DAs with single units that are activating each other and activating themselves (DASA) (**Fig S2 D-F)**.

### Connecting bistable motifs through feedback loops

Next, we probed the effects of adding self-activations to the nodes of the coupled networks. The most frequent states seen in C7 (circuit C2 together with self-activation added on all four nodes) were the same as those seen in C2 – 1001 and 0101, for both RACIPE (∼34% each) and Boolean (∼25% each) simulations (JSD = 0.07; **Fig 3A**). As with a simple toggle switch [12], adding self-activations in Boolean simulations gave rise to states that are not compliant with the behavior of TS motifs (1000, 0001, 1110, 0111, 1100, 0011 – each of these states disagree with at least one of the two toggle switches) with ∼6% frequency each. We refer to such states as ‘hybrid’ from here on. The corresponding STG (**Fig S3A**) reveals interesting dynamics. Out of the 16 possible states for the network, 10 of them behaved as stable states, with 8 of them having a single-state attractor basin. The other six converged to the two most frequent stable states: 1001 and 0110. Interestingly, RACIPE also showed an increase in the frequency of the six hybrid states by 2-4% (C7 vs. C2 frequency distributions), a characteristic that was not seen in TSSA (compare the frequency of 11 and 00 in **Fig 1E** vs. that in **Fig 1A**).

**Fig 3.**
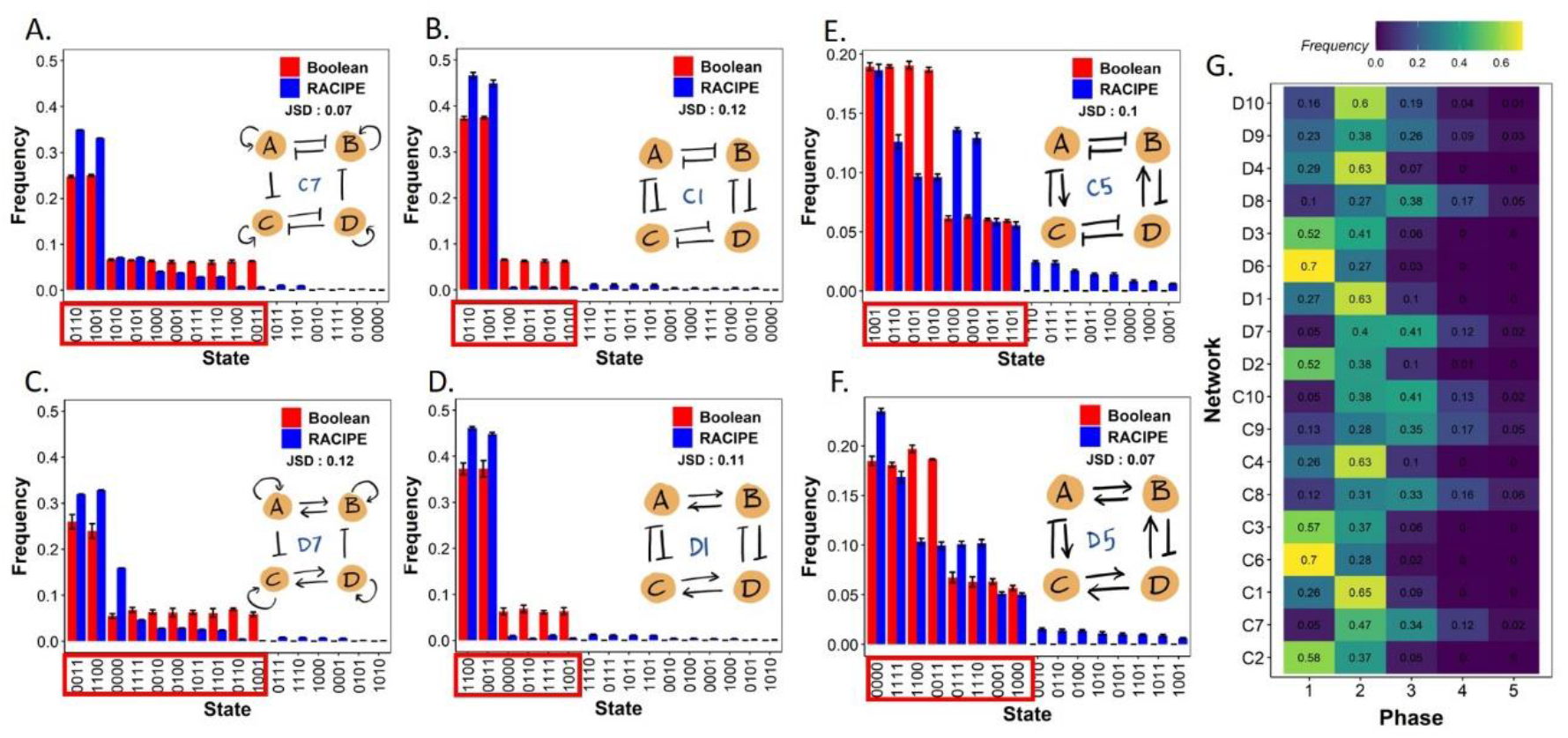
Self-activations and feedback loop couplings. Steady state distribution of A) C7, B) C1, C) C5, D) D7, E) D1, and F) D5. states reported by Boolean simulation are highlighted in the red box. G) Heatmap depicting the frequencies of occurrence of different mono and multistable steady state phases in phenotypic distributions of all 20 (C1-C10, D1-D10) networks.

The addition of self-activations to the C3 network (i.e., C8 network) also gave similar results as those seen in C7 vs. C2 as mentioned above. First, in Boolean simulations, a total of 10 steady states were seen (**Fig S3B**), with 8 of them having a single-state attractor basin (**Fig S3C**). Second, 6 of these are hybrid (i.e., inconsistent with the predominant behavior of individual TS motifs) in Boolean simulations. Third, in RACIPE, an increase of 2-4% in the frequency of hybrid states was observed as compared to that seen in C3. These observations reveal the design principles of adding self-activations on coupled toggle switches through a total of two coupling edges (A activates or inhibits C, and D activates or inhibits B).

While the self-activations are single-edge feedback loops added to the network, one can add feedback loops to the nodes mediated by other nodes. For example, consider network C1 (**Fig 3B**). In this, two toggle switches are connected via double inhibition feedback loops (A and C inhibit each other, and so do B and D). Due to these feedback loops, the net effect on each node is a positive feedback loop mediated via another node (instead of a direct self-activation). Furthermore, similar to a simple inhibition link across the bistable motifs (TA or DS), the double inhibition also imparts an antagonistic behavior between the two nodes that it connects. Thus, intuitively, we expected C1 to act in a similar manner as C7 (which has this positive feedback loop in the form of direct self-activation). Upon comparing the phenotypic distributions of C7 (**Fig 3A**) and that of C1 (**Fig 3B**), we observed in Boolean simulations that both C1 and C7 allow for hybrid states, each of which has a single-state attractor basin (**Fig S3D**). However, while C1 allows for only two hybrid states (1100 and 0011), C7 allows for four additional ones as well (1000, 0111, 1110, 0001). Each hybrid state has a 6.25% frequency; thus, the net frequency of hybrid states in C7 is three-fold that in C1.

Furthermore, the states 1010 and 0101 that we observed in C2 were witnessed in C1 too, but their frequency was approximately one-third of what was seen in C2 (∼17% vs. ∼6%, **Table S1**). Put together, these observations suggest that coupling through double inhibition (i.e., A and C inhibiting each other and similarly B and D inhibiting each other) weakly exhibits the properties of a self-activation (fewer hybrid states seen in C1 vs. those seen in C7) but display a stronger antagonism between the two toggle switches as compared to a simple inhibition (less frequency of 1010 and 0101 – the states that do not obey the antagonistic coupling between the 2 TS motifs – seen in C1 as compared to that in C2). This stronger antagonism also underlies a higher frequency of two predominant states – 0110 and 1001 – in C1 as compared to C2, while comparing RACIPE outputs (compare first two columns in **Fig 3B** vs. that in **Fig 2A**). The differences observed in absolute frequencies of states when using RACIPE vs. Boolean approaches could be attributed to the expected probabilities of a link being “on”, as seen earlier for C2, C3, D2, and D3 (**Table S5**), highlighting common principles.

The trends discussed above while comparing C1, C2, and C7 circuits are also conserved while comparing the frequency distributions for D1, D2, and D7. First, D1 allows for two hybrid states (0110, 1001), each with ∼6.25% frequency, while D7 enables four additional states (0010, 1101, 0100, 1011) too (**Fig 3C-D, Table S3**). Second, each of these six hybrid states has a single-state attractor basin (**Fig S3E, F**). Third, mutual antagonism mediated by mutual inhibition instead of an individual inhibitory link was stronger, as seen via both Boolean and RACIPE outputs. In Boolean, the states 0000 and 1111 were three-fold less frequent in D1 as compared to D2 (compare columns 3 and 5 in **Fig 3D** with columns 3 and 4 in **Fig 2D**). In RACIPE, the frequency of 1100 and 0011 states was higher in D1 as compared to D1 (compare the first two columns in Fig 3D vs. those in Fig 2D). This similarity in comparative analysis for D1, D2, and D7 as those seen in C1, C2, and C7 further endorses that connecting bistable motifs – TS or DA – through mutual inhibition scenarios can drive a strong coupling between the individual bistable motifs while still allowing for certain hybrid states to exist. Similar trends were observed in comparisons of C3, C4, and C8 (C3: A activates C and D activates B; C4: A and C activate each other, and so do B and D; C8: C3 + self-activation for all nodes) (**Table S2, Fig S3G, C**) and for D3, D4 and D8 (same as C3, C4, and C8 but for DA motifs instead of TS motifs) (**Table S4, Fig S3H, I**) reveal that bidirectional mutual activation between bistable motifs drives stronger coordination in the dynamical state space of the two motifs. The strength of these feedback loops as antagonistic/ activating links is also proven by the fact that the addition of self-activations to these networks (C6, C9, D6, D9) doesn’t change the steady state frequency distribution or the state transition graphs (**Fig S4A, B, D, E, G, H, J, K**).

As a control case, we next coupled the bistable motifs (TS, DA) through negative feedback loops (C5: C inhibits A and A activates C, similarly; B inhibits D and D activates B; D5: same as C5 but for DA motifs instead of TS ones). This coupling does not lead to any predominant states as seen earlier in coupling through C1-C4 or D1-D4 networks (**Fig 3E; S3J, K**), further endorsing the design principles of certain types of coupling between the bistable motifs. C5 and D5 did, however, have a property similar to C1, C4, D1, and D4, in that the addition of self-activation does not change the Boolean phenotypic distribution and the state transition graph much (**Fig S4 C, F, I, L**). The RACIPE results of D10 do show an increase in the frequency of the state 0000.

### Effect of coupling on multistability of bistable motifs

So far, we have analyzed the effect of the coupling of bistable motifs on phenotypic distribution. We next wanted to understand the signatures of multistability that these couplings can result in. To begin with, we computed the frequencies of multistable phases in all networks considered (**Fig 3G**). As seen earlier with TS and DA **(Fig S1B, D)**, the addition of self-activation invariably created phases with more than two stable states (columns 3,4 and 5 in Fig 3G). Furthermore, in networks not involving direct self-activation on any nodes (C1-C5, D1-D5), the frequency of bistable phases (column 2) was higher for the connections mediated by positive feedback loops (C1, C4, D1, D4) as opposed to those by negative feedback loops (C5, D5) (**Fig 3G**).

We next probed into the constitution of multistable phases seen in coupled bistable motifs. We started with circuits having single inhibition or activation coupling (C2, C3, D2, D3). A single toggle switch was able to give rise to 2 states – 01, 10 – and one bistable phase {01, 10} (**Fig 1D)**. Coupling two toggle switches with single activation/inhibition links (C2, C3) maintained this trend, i.e., bistable phases comprising of the two most dominant states (**Fig 4A**) – {0101, 1010} for C3 and {1001, 0110} for C2. Similarly, the most dominant monostable phases were also comprised of the two most dominant states in the steady state frequency(**FigS5A-B**). C2 and C3 circuits also enabled tristable phases but with relatively low frequency (<10%; **Fig 3G**). Phases consisting of 4 or more co-existing states have frequencies less than 0.01% and can therefore be neglected. The net frequency of bistable phases seen in C2 and C3 is still less than that of monostable phases, approximately by the same fraction (∼0.5) as seen in a TS. Reinforcing patterns showed up in D2, D3 circuits in terms of frequency and composition of multistable phases (**Fig 3G, 4B, Fig S5A-B**).

**Fig 4.**
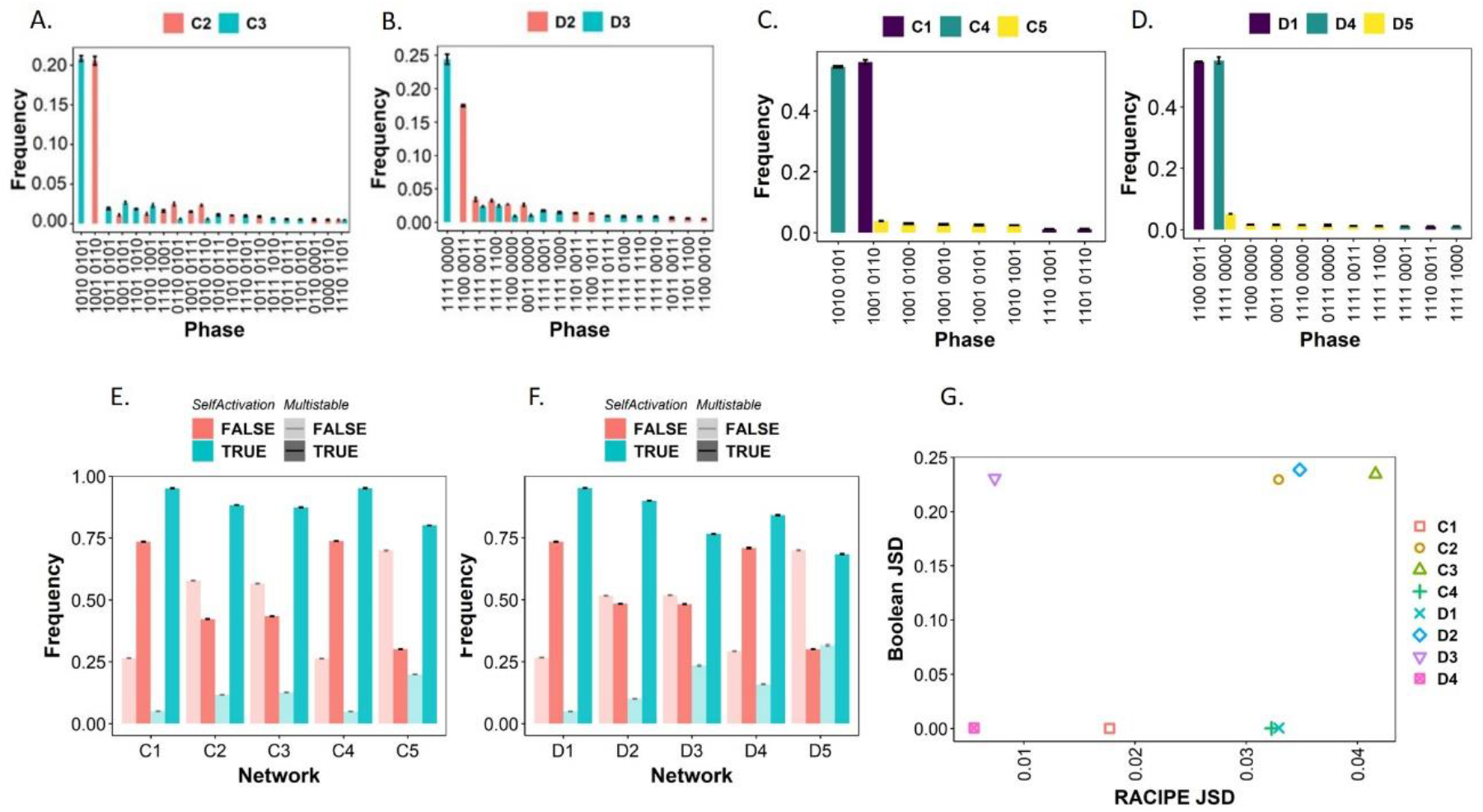
Effect of couplings on multistability. Bistable phase distribution for the networks A) C2 and C3. B) D2 and D3. C) C1, C4, and C5 and D) D1, D4, and D5 E) Comparison of monostable (transparent bars) and multistable (opaque bars) frequencies of networks C1-6 with (green) and without (red) self-activations. F) Same as E but for D1-D5. G) Scatterplot of RACIPE and Boolean JSD values of all the networks.

Next, we investigated the behavior of circuits that had direct mutual inhibition or activation between two nodes belonging to two different bistable motifs being coupled (C1, C4, D1, D4). These networks showed a similar composition of multistable phases as seen in C2, C3, D2, and D3 circuits; however, the frequency of bistable phases increased approximately two-fold (compare C1 vs. C2, D1 vs. D2, C3 vs. C4, D3 vs. D4) (**Fig 4 C-D, Fig S5 E-H**). This increase was largely at the expense of decreased frequency of monostable phases; this trend seems to emerge from a larger number of positive feedback loops embedded in the networks C1, C4, D1, and D4 when compared to C2, C3, D2, and D3 correspondingly [7].

The phases in C5 and D5 were largely monostable (∼70%), with a smaller fraction of bistable phases (∼27%). The frequency order of the monostable phases (**Fig S5C-D**) remained the same as that of the states in phenotypic distribution (**Fig 3E, H**). However, the lack of predominant states, as seen in C1-C4 and D1-D4 networks, influenced the composition of bistable phases as well. Consequently, unlike C1-C4 and D1-D4 networks, C5 and D5 networks did not exhibit any predominant bistable phase; each such phase instead had only ∼2% frequency (**Fig 4C-D**).

The addition of self-activation to the nodes of these networks had little effect on the composition of various phases (**Fig S6 A-H**) but increased the frequency of tristable phases in all cases (Compare **Fig S5E-H** with **Fig S6I-L**). One anomaly being observed was the dominance of 0000 state as one of the constituents in multistable phases seen in D4, D8, and D9 (**Fig SD, H; S6D, H, L**), suggesting that when DA switches are connected only by activation links, the addition of direct or indirect self-activation leads to the enhancement of the occurrence of 0000 state in the outcome. The weaker this extent of self-activation, the less likely is the expected enrichment of 0000 state, a case made by D3 (**Fig 4B; S5B, F**).

Generally, however, the addition of self-activations increased the frequency of multistable phases but decreased that of monostable phases (**Fig 4E-F**). Despite the change in a phase frequency distribution, phenotypic distribution of the networks does not always change upon adding self-activation (**Fig 4G**), quantified by JSD (information difference) [30] between phenotypic frequencies of networks with and without self-activations. RACIPE JSD is very low, whereas the Boolean JSD is also low for six networks and slightly high for four networks: C2, D2, C3, and D3. This difference can be explained by visualizing that simple inhibition/activation links apply weaker restrictions in the coupled state space as compared to the feedback loops and thus are more susceptible to imparting hybrid phenotypes upon adding self-activation.

## Discussion

Network motifs, defined as patterns of regulatory connections that frequently occur in many biological networks, are viewed as the building blocks of complex regulatory topologies [35,36]. While the behavior of various network motifs has been well understood individually, the question of how does coupling or interlinking such regulatory motifs govern their behavior is relatively less understood [37]. Here, we take a step in this direction, trying to understand the design principles that govern the behaviors of coupled bistable switches.

First, we investigate how do different forms of coupling affect the phenotypic distribution of bistable switches coupled. Second, we also interrogate how do such couplings influence multistability, i.e., how is the ability for multiple phenotypes to co-exist, and more importantly, the ability for a population to regain its phenotypic distribution modulated upon such coupling. We used two different approaches to analyze the coupled networks: a continuous formalism of network simulation (RACIPE) and a discrete one (Boolean) inspired by the Ising model of ferromagnetism [38]. The Boolean formalism allowed us to understand the effect of network topology alone on the dynamical behavior, while RACIPE also enabled investigating the behavior over a wide ensemble of kinetic parameters in coupled ODEs. We explored 20 different coupled networks in total, out of which 16 of them were synergistic, i.e., no two nodes in the networks are connected, directly or indirectly, via conflicting links which can add ‘frustration’ to the states. C5, D5, C10, and D10 are the four ‘non-synergistic’ networks where the node pairs {A, C} and {B, D} are connected by both inhibiting and activating links, thus leading to high frustration in states [39] (**Fig 5)**.

**Fig 5.**
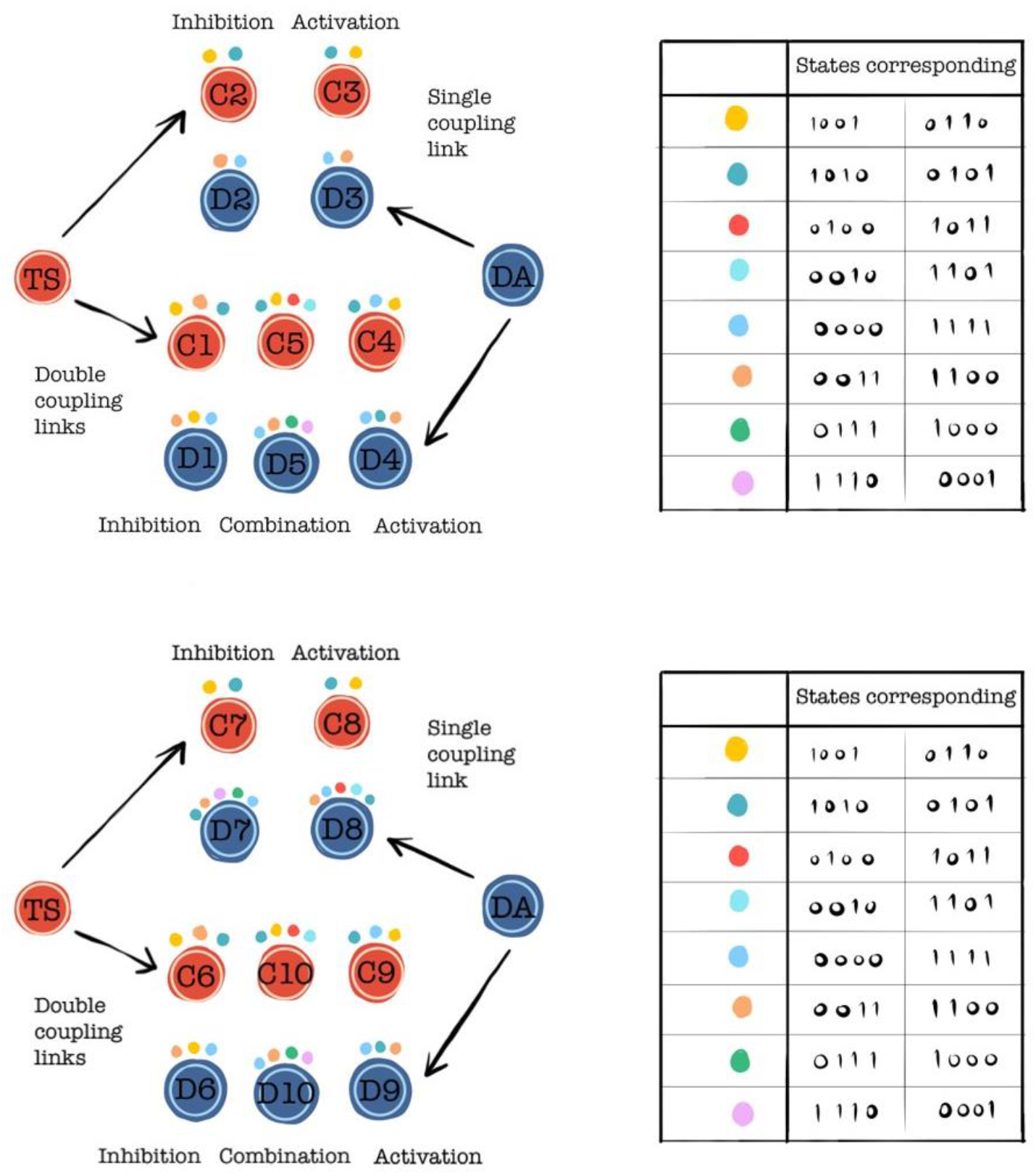
Summary of the behavior of synergistic coupling between bistable switches.

For networks without self-activation, phenotypic distributions intuitively followed the “rules” of the network topology. The states that followed the network topology fully had the highest frequency, and the frequency kept decreasing as the compliance with topology decreased. When bistable switches are connected with inhibitions or activations (C2, C3, D2, D3), both Boolean and RACIPE results suggested that the most predominant states obtained cannot oppose the rules of individual bistable motifs (both nodes in a TS have opposite values – 10 or 01, and those in a DA have the same value – 00 or 11). Thus, the occurrence of hybrid states with respect to individual bistable switches was not observed. However, when the motifs relate to strong positive feedback loops (C1, C4, D1, D4), we see hybrid states emerging. One way to interpret this trend is that bistable switches (TS and DA) are dominant over their single link counterparts (Inhibition and activation respectively; C2, C3, D2, D3) but not over feedback loops formed across them (C1, C4, D1, D4). In other words, adding feedback loops as seen in C1, C4, D1, and D4 can be considered akin to adding self-activation to the coupled networks, thus enabling hybrid states as seen for bistable switches [12]. This interpretation also helps in understanding how adding direct self-activation on individual nodes (C7, C8, D7, D8) facilitates various hybrid phenotypes in the coupled system.

Given the two possible steady states for each bistable switch (10 and 01 for TS, 00 and 11 for DA), in the case of weakly coupled cases, we expect to have four possible states. Interestingly, the couplings studied here restrict the number of possible phenotypes. While single inhibition/ activation couplings only reduce the frequency of some phenotypes, networks connected with positive feedback loops can reduce the number of phenotypes as well. Similar observations can be made for couplings of bistable motifs with self-activation.

Unlike phenotypic composition, multistability has straightforward patterns. The addition of self-activation to the networks always increases multistability, with the most dominant phases always composed of the most dominant phenotypes observed in the phenotypic frequency. One important aspect to note when comparing multistability and phenotypic composition is that we observed an increase in the frequency of multistability upon adding self-activation, but the corresponding change in the phenotypic distributions, as measured by JSD, was very less. This result is similar to our previous observations [7], where a change in multistability frequency did not correlate with a change in phenotypic distribution. Biologically, the phenotypic distribution represents a snapshot of a population, while multistability depicts the dynamic evolution of that snapshot (i.e., ability to switch back and forth among multiple states enabled). Hence, the observation that these two properties do not necessarily correlate is not a surprise. However, many of the biological networks are derived from such snapshot data, and hence, one must be careful while exploring the dynamics of such networks.

We should mention this exercise is not a comprehensive analysis of different coupling of bistable switches but can serve as a first step towards understanding the nature of these couplings based on the changes in phenotypic distributions and multistability patterns. For instance, we have not considered coupling between a TS and a DA. Nonetheless, we were able to decode a set of design principles of synergistic couplings between bistable switches.

## Author contributions

M.K.J and K.H. designed the research; K.H., P.H., A.G., and V.U. carried out the simulations; K.H., P.H., and A.S.D. analyzed the data; all authors discussed results and participated in the preparation of the paper; M.K.J. supervised the research.

