## Supplemental figures and tables for "Emergent properties of coupled bistable switches"

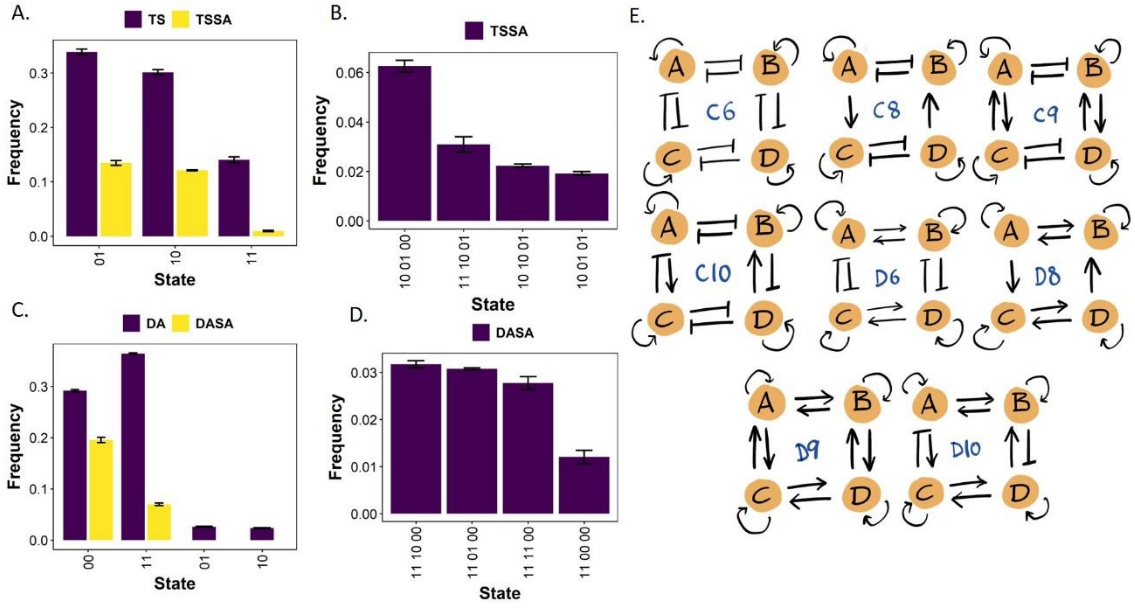


**FigS1.** A) Comparison of monostable steady state distribution of TS and TSSA. B) Tristable steady state distribution of TSSA. C) Comparison of monostable steady state distribution of DA and DASA. D) Tristable steady state distribution of DASA. E) Different variations of coupling between two TS or DA that were evaluated in the study.

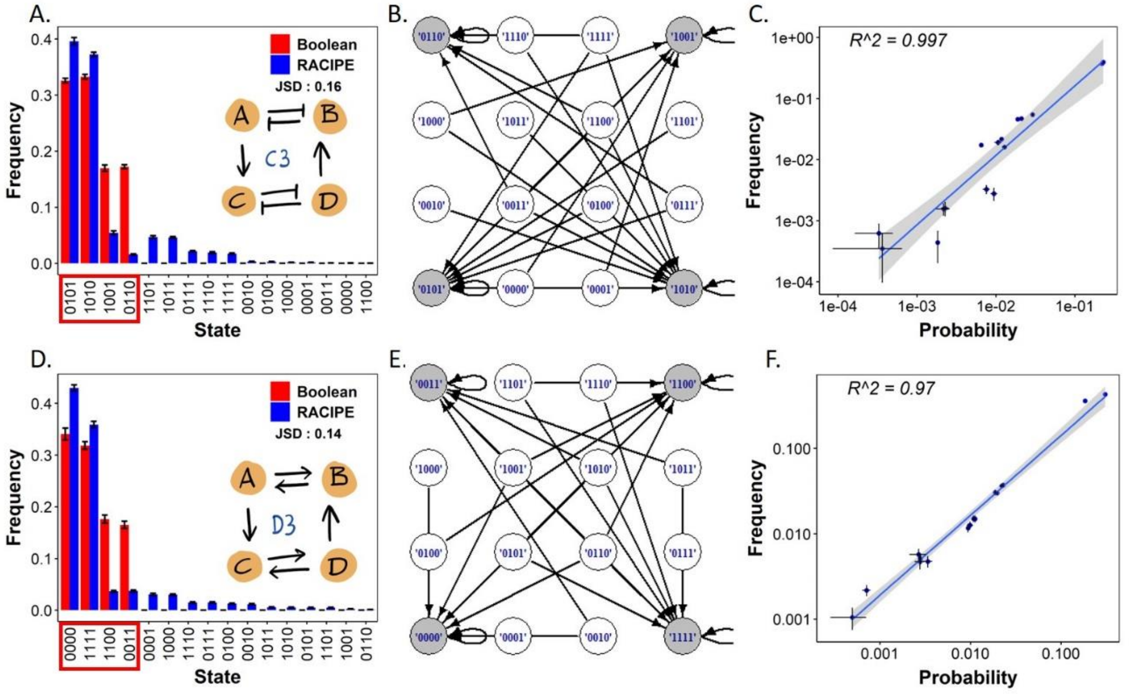


**FigS2.** A) Phenotypic distribution of C3. B) State transition graph of C3. C) Link-strength analysis of parameter sets of C3. D) Steady state distribution of D3. E) State transition graph of D3. F) Link-strength analysis of parameter sets of D3. In each phenotypic distribution, states occurring in Boolean simulations are highlighted in red box.

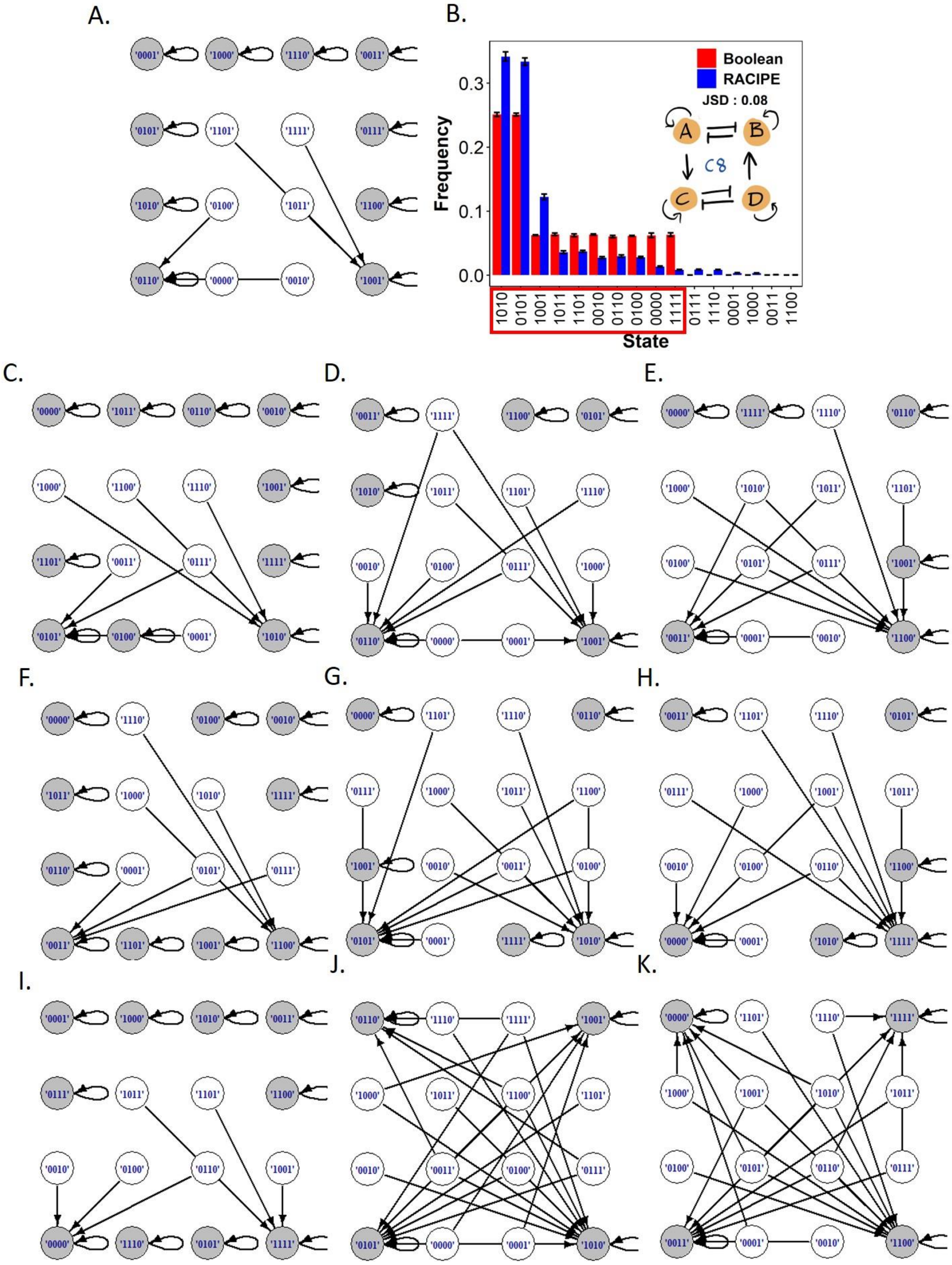


**FigS3.** A) State transition graph of C7. B) Steady state distribution of C8. C) State transition graphs of C) C8, D) C1, E) D1, F) D7, G) C4, H) D4, I) D8, J) C5 and K) D5.

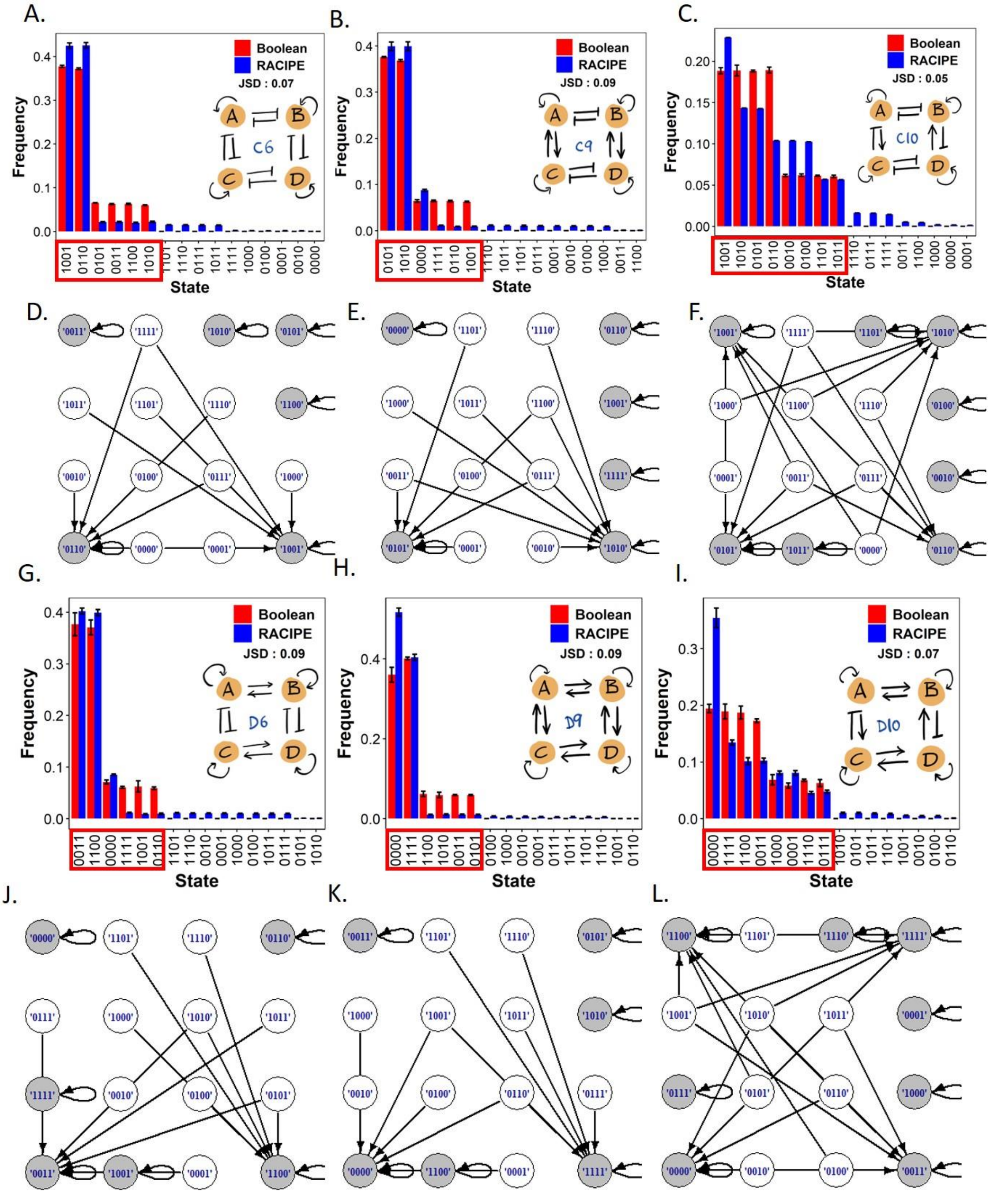


**FigS4.** Steady state distributions of A) C6, B) C9, C) C10, G) D6, H) D9 and I) D10. State transition graphs of D) C6, E) C9, F) C10, J) D6, J) D9 and L) D10.

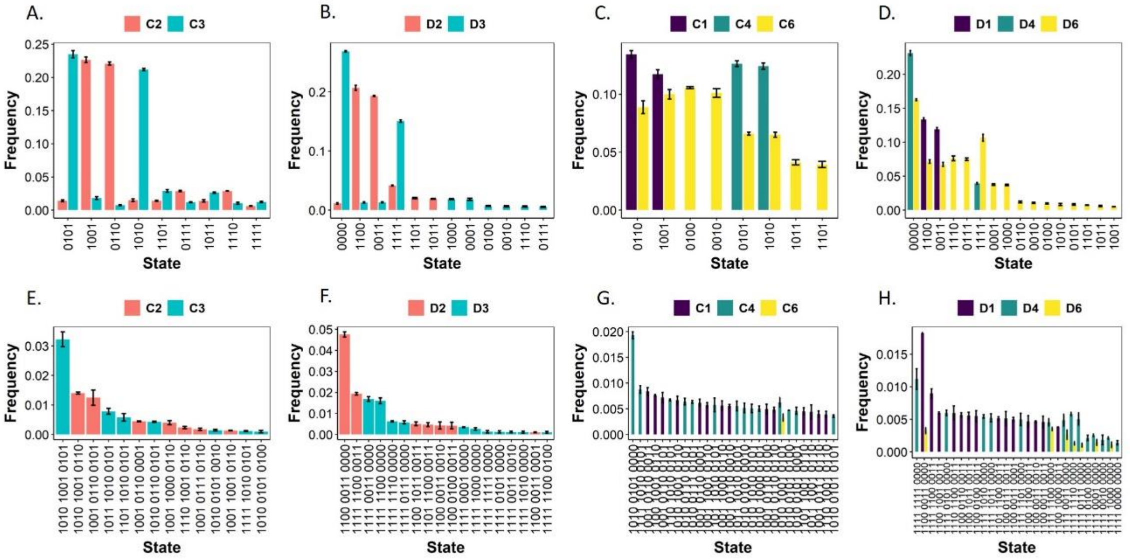


**FigS5.** A) Comparison of monostable steady state distributions of C2 and C3. B) Comparison of monostable steady state distributions of D2 and D3. C) Comparison of monostable steady state distributions of C1, C4 and C6. D) Comparison of monostable steady state distributions of D1, D4 and D6. E) Tristable steady state distributions of networks C2 and C3. F) Tristable steady state distributions of networks D2 and D3. G) Comparison of tristable steady state distributions of C1, C4 and C6. H) Comparison of tristable steady state distributions of D1, D4 and D6.

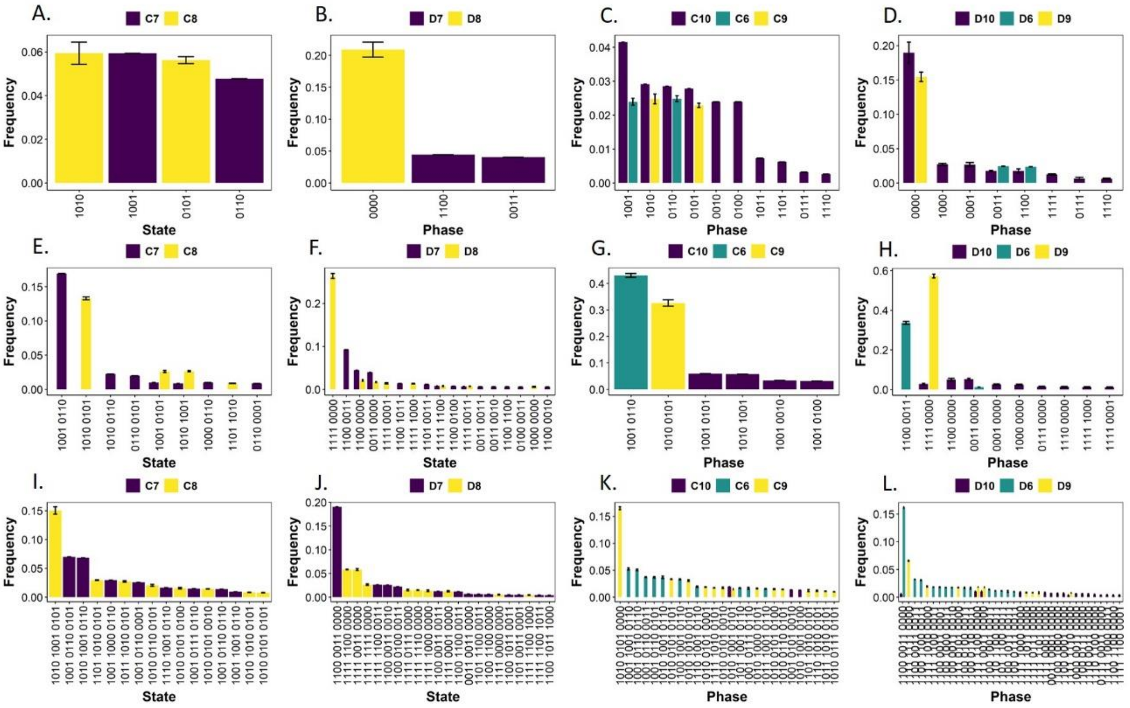


**FigS6.** A) Monostable phase distribution of A) C8 and C9, B) D8 and D9, C) C10, C12 and C7 and D) D10, D12 and D7. E-H) Same as A-D for Bistable phase distribution. I-L) Same as A-D but for tristable phase distribution.

**Table S1: Comparison of steady state frequencies of C1, C2 and C7 (mean across 3 replicates)**

| State | Boolean | | | RACIPE | | |
|---|---|---|---|---|---|---|
| | C1 | C2 | C7 | C1 | C2 | C7 |
| '0110' | 0.373 | 0.33 | 0.247 | 0.466 | 0.381 | 0.349 |
| '1001' | 0.374 | 0.325 | 0.25 | 0.449 | 0.379 | 0.331 |
| '1110' | 0 | 0 | 0.06 | 0.012 | 0.043 | 0.029 |
| '0111' | 0 | 0 | 0.061 | 0.011 | 0.042 | 0.029 |
| '1011' | 0 | 0 | 0 | 0.011 | 0.022 | 0.011 |
| '1101' | 0 | 0 | 0 | 0.011 | 0.023 | 0.01 |
| '0011' | 0.063 | 0 | 0.063 | 0.006 | 0.002 | 0.007 |
| '1100' | 0.066 | 0 | 0.062 | 0.006 | 0.002 | 0.008 |
| '1010' | 0.062 | 0.172 | 0.066 | 0.005 | 0.039 | 0.071 |
| '0101' | 0.062 | 0.174 | 0.065 | 0.005 | 0.037 | 0.071 |
| '0001' | 0 | 0 | 0.061 | 0.004 | 0.009 | 0.038 |
| '1000' | 0 | 0 | 0.063 | 0.004 | 0.009 | 0.041 |
| '1111' | 0 | 0 | 0 | 0.004 | 0.008 | 0.002 |
| '0100' | 0 | 0 | 0 | 0.004 | 0.001 | 0.002 |
| '0010' | 0 | 0 | 0 | 0.004 | 0.001 | 0.002 |
| '0000' | 0 | 0 | 0 | 0 | 0 | 0 |

**Table S2: Comparison of steady state frequencies of C3, C4 and C8 (mean across 3 replicates)**

| State | Boolean | | | RACIPE | | |
|---|---|---|---|---|---|---|
| | C3 | C4 | C8 | C3 | C4 | C8 |
| '0101' | 0.326 | 0.374 | 0.251 | 0.396 | 0.454 | 0.333 |
| '1010' | 0.333 | 0.375 | 0.251 | 0.372 | 0.454 | 0.341 |
| '1001' | 0.169 | 0.065 | 0.062 | 0.054 | 0.004 | 0.122 |
| '1101' | 0 | 0 | 0.062 | 0.047 | 0.011 | 0.037 |
| '1011' | 0 | 0 | 0.064 | 0.046 | 0.012 | 0.036 |
| '0111' | 0 | 0 | 0 | 0.022 | 0.012 | 0.008 |
| '1110' | 0 | 0 | 0 | 0.019 | 0.011 | 0.008 |
| '1111' | 0 | 0.062 | 0.063 | 0.017 | 0.01 | 0.008 |
| '0110' | 0.172 | 0.063 | 0.06 | 0.016 | 0.005 | 0.03 |
| '0010' | 0 | 0 | 0.063 | 0.003 | 0.004 | 0.027 |
| '0100' | 0 | 0 | 0.061 | 0.003 | 0.004 | 0.028 |
| '1000' | 0 | 0 | 0 | 0.002 | 0.004 | 0.003 |
| '0001' | 0 | 0 | 0 | 0.002 | 0.004 | 0.003 |
| '0011' | 0 | 0 | 0 | 0.001 | 0.001 | 0.001 |
| '0000' | 0 | 0.062 | 0.062 | 0 | 0.011 | 0.013 |
| '1100' | 0 | 0 | 0 | 0 | 0.001 | 0 |

**Table S3: Comparison of steady state frequencies of D1, D2 and D7 (mean across 3 replicates)**

| State | Boolean | | | RACIPE | | |
|---|---|---|---|---|---|---|
| | D1 | D2 | D7 | D1 | D2 | D7 |
| '1100' | 0.372 | 0.32 | 0.24 | 0.461 | 0.375 | 0.328 |
| '0011' | 0.373 | 0.327 | 0.26 | 0.448 | 0.362 | 0.32 |
| '1110' | 0 | 0 | 0 | 0.012 | 0.009 | 0.008 |
| '1111' | 0.061 | 0.172 | 0.068 | 0.011 | 0.085 | 0.047 |
| '0111' | 0 | 0 | 0 | 0.011 | 0.009 | 0.009 |
| '1011' | 0 | 0 | 0.063 | 0.011 | 0.033 | 0.026 |
| '1101' | 0 | 0 | 0.061 | 0.011 | 0.034 | 0.025 |
| '0000' | 0.063 | 0.181 | 0.055 | 0.01 | 0.055 | 0.159 |
| '1001' | 0.063 | 0 | 0.059 | 0.005 | 0.003 | 0.001 |
| '0010' | 0 | 0 | 0.063 | 0.004 | 0.01 | 0.028 |
| '0100' | 0 | 0 | 0.062 | 0.004 | 0.008 | 0.029 |
| '0110' | 0.068 | 0 | 0.07 | 0.004 | 0.001 | 0.004 |
| '0001' | 0 | 0 | 0 | 0.003 | 0.007 | 0.006 |
| '1000' | 0 | 0 | 0 | 0.003 | 0.007 | 0.007 |
| '0101' | 0 | 0 | 0 | 0.001 | 0.001 | 0.001 |
| '1010' | 0 | 0 | 0 | 0.001 | 0.001 | 0.001 |

**Table S4: Comparison of steady state frequencies of D3, D4 and D8 (mean across 3 replicates)**

| State | Boolean | | | RACIPE | | |
|---|---|---|---|---|---|---|
| | D3 | D4 | D8 | D3 | D4 | D8 |
| '0000' | 0.341 | 0.369 | 0.263 | 0.43 | 0.534 | 0.474 |
| '1111' | 0.318 | 0.387 | 0.252 | 0.359 | 0.382 | 0.303 |
| '0011' | 0.165 | 0.066 | 0.057 | 0.037 | 0.005 | 0.051 |
| '1100' | 0.176 | 0.057 | 0.062 | 0.036 | 0.005 | 0.054 |
| '0001' | 0 | 0 | 0.06 | 0.03 | 0.011 | 0.03 |
| '1000' | 0 | 0 | 0.054 | 0.03 | 0.011 | 0.03 |
| '1110' | 0 | 0 | 0.062 | 0.015 | 0.005 | 0.016 |
| '0111' | 0 | 0 | 0.062 | 0.015 | 0.004 | 0.016 |
| '0100' | 0 | 0 | 0 | 0.013 | 0.01 | 0.005 |
| '0010' | 0 | 0 | 0 | 0.012 | 0.011 | 0.006 |
| '1011' | 0 | 0 | 0 | 0.006 | 0.005 | 0.004 |
| '1010' | 0 | 0.065 | 0.064 | 0.005 | 0.005 | 0.004 |
| '0101' | 0 | 0.057 | 0.064 | 0.005 | 0.006 | 0.004 |
| '1101' | 0 | 0 | 0 | 0.005 | 0.005 | 0.004 |
| '1001' | 0 | 0 | 0 | 0.002 | 0.001 | 0.001 |

| | | | | | | |
|---|---|---|---|---|---|---|
| '0110' | 0 | 0 | 0 | 0.001 | 0 | 0 |

**Table S5: Linear fit parameters between state probability calculated from RACIPE parameter sets and RACIPE steady state frequency**

| Network | Slope | Intercept | R-squared |
|---|---|---|---|
| C1 | 2.2 | -0.0011 | 1 |
| C2 | 1.8 | -0.00041 | 0.973 |
| C3 | 1.7 | -0.00011 | 0.997 |
| C4 | 2 | -0.00087 | 1 |
| C5 | 1.8 | 0.009 | 0.881 |
| C6 | 2.5 | -0.0051 | 0.997 |
| C7 | 2.2 | -0.0054 | 0.995 |
| C8 | 2 | -0.0032 | 0.997 |
| C9 | 2.8 | -0.0042 | 0.987 |
| C10 | 2.2 | 0.0039 | 0.982 |
| D1 | 2 | -0.001 | 1 |
| D2 | 1 | -0.00076 | 0.998 |
| D3 | 1.5 | 0.0023 | 0.97 |
| D4 | 1.8 | 0.001 | 0.992 |
| D5 | 1.5 | 0.014 | 0.942 |
| D6 | 2.3 | -0.0038 | 0.989 |
| D7 | 1.9 | -0.002 | 0.98 |
| D8 | 1.7 | 0.0038 | 0.99 |
| D9 | 2 | 0.0027 | 0.992 |
| D10 | 1.8 | 0.0084 | 0.967 |
